# SpaMOAL: A spatial multi-omics graph contrastive learning method for spatial domains identification

**DOI:** 10.64898/2026.02.23.707433

**Authors:** Jinxia Wang, Yuying Huo, Rui Zhao, Yan Pan, Han Wang, Xiangyu Li

## Abstract

Recent advances in spatial multi-omics technologies have opened new avenues for characterizing tissue architecture and function *in situ*, by simultaneously providing multimodal and complementary information—such as spatially resolved transcriptomic, epigenomic, and proteomic features. Current computational approaches face substantial challenges such as effective integration of multi-omics molecular information with spatial information and corresponding high-resolution histology images. To address this challenge, we proposed SpaMOAL (**Spa**tially **M**ulti-**O**mics graph contr**A**stive **L**earning), a graph-based contrastive learning approach for spatial domain identification. SpaMOAL learns clustering-friendly representations from spatial multi-omics data by integrating spatial coordinates, histological image features and molecular profiles, enabling accurate delineation of spatial tissue domains. Benchmarking across multiple recent paired spatial multi-omics datasets demonstrated that SpaMOAL consistently outperforms existing methods. By enabling accurate spatial domain delineation, SpaMOAL provides a powerful framework for interpreting tissue organization and cellular microenvironments.

## Introduction

Spatial multi-omics technologies have emerged as a powerful approach for comprehensively dissecting complex tissues, providing insights beyond the reach of single-omics methods. Recent advances have enabled simultaneous profiling of multiple molecular features with spatial resolution. These technologies can be broadly categorized into sequencing-based and imaging-based strategies, each offering distinct advantages in resolution, throughput, and molecular coverage. Sequencing-based methods include DBiT-seq^1^, spatial-CITE-seq^2^, spatial ATAC-RNA-seq (assay for transposase-accessible chromatin and RNA using sequencing), CUT&Tag-RNA-seq^3^, SPOTS^4^, SM-Omics^5^, Stereo-CITE-seq^6^, spatial RNA-TCR-seq^7^, and 10x Genomics Xenium Protein^8^. These methods rely on spatially barcoded molecular capture combined with next-generation sequencing to quantify multiple omics layers. In contrast, imaging-based platforms, including seqFISH+^9^, MERFISH^10^, MERSCOPE for DNA^11^ and NanoString CosMx^12^, directly visualize molecular targets in situ through high-throughput multiplexed fluorescence imaging, providing subcellular spatial resolution. These technologies typically produce two or more spatially resolved omics layers from the same tissue section, often accompanied by corresponding high-resolution hematoxylin and eosin (H&E) stained images.

Effective integration of complementary data modalities provides a multidimensional view of tissue architecture and function, thereby enhancing our understanding of biological processes and disease mechanisms. Integrating heterogeneous modalities into unified low-dimensional representations is central to spatial multi-omics analysis, enabling downstream analysis such as domain detection. However, it is challenged by the distinct feature spaces of different modalities. Converting them into a common feature space based on prior knowledge can mitigate these differences, but often results in information loss. Moreover, most existing methods suffer from not fully leveraging spatial information modeling and matched high-resolution histological images. For example, traditional single-cell multi-omics integration methods such as Seurat WNN^13^, MOFA+^14^, and totalVI^15^ cannot incorporate spatial coordinate information. Some spatial transcriptomics clustering approaches, like MUSE^16^, SpaGCN^17^, and SpaDAC^18^, can effectively integrate transcriptomics, spatial location, and morphological imaging data to identify spatial domains. However, these approaches are limited as they cannot incorporate additional omics modalities beyond transcriptomics. Most recent methods like SpatialGlue^19^ and MISO^20^ aim to learn unified low-dimensional representations from spatial multi-omics data. However, SpatialGlue does not support histology imaging as input, while MISO lacks explicit modeling of spatial coordinates. Additionally, MISO requires manual specification of low-quality modalities based on biological knowledge to mitigate their negative effects.

To overcome all the challenges mentioned above, we present a spatially multi-omics graph contrastive learning method (SpaMOAL) to identify spatial domains. SpaMOAL integrates molecular, spatial and morphological information in a unified embedding space. Specifically, it aligns cross-modal shared representations while preserving modality-specific components, thereby enabling coherent integration of heterogeneous modalities within a unified latent space. The concatenated embedding of the shared and private representations effectively combines complementary information, yielding a robust multimodal representation. Benchmarking on both simulated and real-world datasets demonstrates that SpaMOAL consistently achieves superior performance compared to state-of-the-art spatial multi-omics methods.

## Materials and methods

### Overview of SpaMOAL workflow

The basic idea of SpaMOAL is to learn multimodal representations through a graph contrastive autoencoder framework that integrates multi-omics molecular profiles, spatial coordinates and morphological information. It aims to capture both shared and modality-specific variation across modalities within a unified representation space^21^. As illustrated in Fig. 1, SpaMOAL first constructs an adjacency matrix based on spatial coordinates and augments it with omics- or image-derived features to build modality-specific spatial neighbor graphs, where each node represents an individual spot or cell. Modality-specific graph encoders are then employed to obtain low-dimensional representations that are further processed by a multilayer perceptron (MLP) module to separate shared and modality-specific components. To optimize these representations, SpaMOAL jointly leverages four complementary objectives to balance reconstruction-based structural consistency, cross-modal alignment, and modality-specific discrimination. A reconstruction term ensures accurate node-feature reconstruction and preserves neighborhood consistency in each modality’s spatial graph. A correlation loss encourages statistical independence between shared and modality-specific representations by minimizing the Pearson correlation coefficients between them. A matching objective further aligns shared representations across modalities, while a contrastive objective enhances within-modality discrimination by pulling similar spots/cells closer and pushing dissimilar ones apart within spatially local microenvironments in the private embedding space. The learned shared representations and private representations are concatenated into unified representations for downstream analyses, particularly for identifying spatial domains through clustering.

**Fig. 1.**
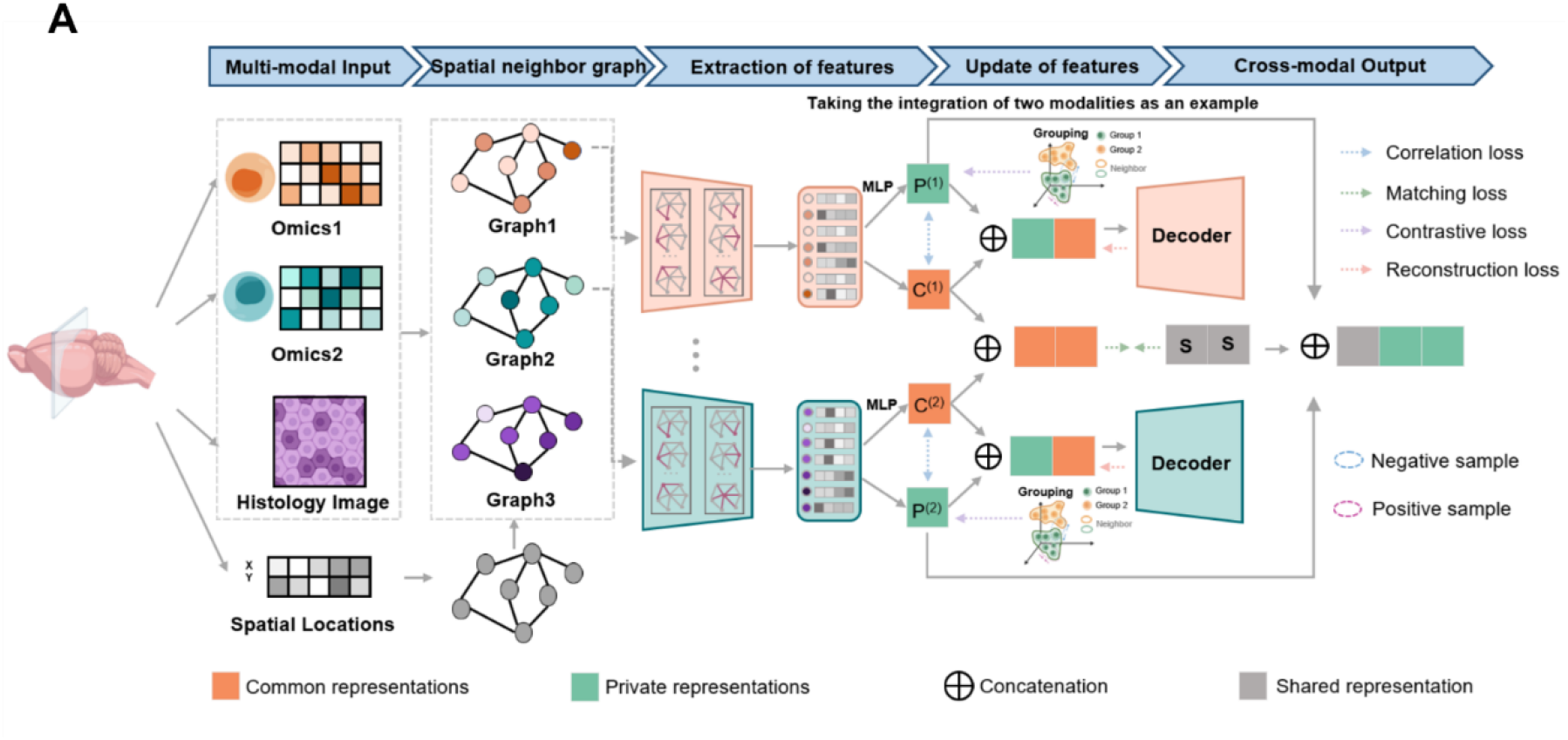
Overview of SpaMOAL. SpaMOAL first uses the k-nearest-neighbor (KNN) algorithm to construct an adjacency matrix based on spatial coordinates and augments it with omics- or image-derived features to build modality-specific spatial neighbor graphs. For each modality, graph encoders are employed to obtain low-dimensional representations that are further processed by a MLP module to separate shared and modality-specific components. To optimize these representations, SpaMOAL jointly leverages four losses to balance reconstruction-based structural consistency, cross-modal alignment, and modality-specific discrimination. The reconstruction loss ensures accurate node-feature reconstruction and preserves neighborhood consistency in each modality’s spatial graph. The correlation loss encourages statistical independence between shared and modality-specific representations by minimizing the Pearson correlation coefficients between them. The matching loss further aligns shared representations across modalities, while a contrastive loss enhances within-modality discrimination among spatial neighbors by pulling similar spots/cells closer and pushing dissimilar ones apart within spatially local microenvironments in the private embedding space. Finally, shared representations and private representations are concatenated into unified representations for downstream analyses.

SpaMOAL is an efficient and scalable algorithm for integrative multi-omics analysis, capable of simultaneously processing spatial transcriptomics, spatial proteomics, spatial chromatin accessibility, and other omics data, together with spatial coordinates and morphological images.

### Algorithm of SpaMOAL

#### Data preprocessing

SpaMOAL provides standardized preprocessing steps for RNA, protein, and chromatin accessibility, while allowing user-defined customization. For transcriptomic data, spots expressing fewer than 10 genes were filtered out. Gene expression counts were then log-transformed and normalized by library size using the SCANPY package, and the top 3,000 highly variable genes (HVGs) were selected for graph construction. For the ADT (antibody-derived tag) data, the counts were normalized using the centered log-ratio across spots, followed by principal component analysis (PCA)^22^. For the chromatin accessibility data, a GeneScore matrix was constructed from peak counts using Seurat, followed by read-count (RC) normalization and selection of the top 3,000 highly variable features through variance-stabilizing transformation. For imaging data, such as H&E-stained histology or immunofluorescence images, SpaMOAL directly processes raw high-resolution images and performs feature extraction using pretrained CNNs.

#### Spatial neighbor graph construction

For each spot, SpaMOAL segments a corresponding image tile centered on its spatial coordinates from the raw histology image. Morphological features are extracted from these tiles using ImageNet pretrained CNNs (Inception_v3^23^ or ResNet50^24^) via transfer learning. All layers except the classifier are retained, and the output of the final convolutional layer is used as the spot-level morphological representation.

Based on the spatial coordinates, SpaMOAL constructs adjacency matrix and builds modality-specific spatial neighbor graphs *G* = {*G*^(1)^, *G*^(2)^, …, *G*^(*R*)^}, where *R* is the number of modalities. Each graph *G*^(*r*)^ = {*V, E*} corresponds to modality *r*, with node set *V* = {*v*_1_, *v*_2_, …, *v*_*N*_} and edge set *E*. Spatial relationships are defined by the Euclidean distances between spots’ spatial coordinates: two spots are connected if one lies among the *k* (default *k* = 6, optional) nearest neighbors of the other.

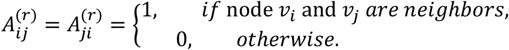

The adjacency matrix *A*^(*r*)^ ∈ *R*^*N*×*N*^, self-loops are added to each spot, and *N* denotes the number of nodes. For each modality, the node features are defined as *X*^(*r*)^ = 𝒯(*X*) ∈ *R*^*N*×*F*^, where 𝒯 denotes the random dropout and *X* denotes the original node features, F denotes the dimension of node features. To ensure dimensional consistency across modalities, *F* is unified to 3,000 (the default target dimension), with lower-dimensional modalities zero-padded to match this size.

#### Graph autoencoder

SpaMOAL first employs a graph convolutional layer *g*^(*r*)^ to each modality to generate node representations *H*^(*r*)^ based on the node features and the adjacency matrix,

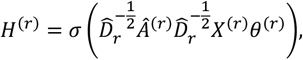

where *Â* ^(^*r*^)^ = *A*^(*r*)^ + *ωI*_*N*_, and *ω* indicates the weight of self-connections. 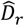 is the degree matrix of *Â* ^(*r*)^, *σ* is the activation function, and *θ*^(*r*)^ is the trainable weight matrix of *g*^(*r*)^. The encoded representations *H*^(*r*)^ are passed through two MLPs with unshared parameters (i.e., 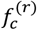 and 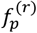) to derive common and private/modality-specific representations (i.e., *C*^(*r*)^ and *P*^(*r*)^), respectively.

1. **The matching loss**. To encourage common representations learnt from different modalities to align toward a unified latent space, SpaMOAL introduces a shared variable *S* as a global reference. All modality-specific common representations [*C*^(1)^, …, *C*^(*R*)^] are concatenated and subjected to singular value decomposition (SVD) to obtain an orthogonality basis that captures the dominant shared structure among modalities. The matching loss then minimizes the distance between each modality’s common representations *C*^(*r*)^ and the shared variable *S*, formulated as:

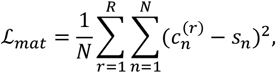

where 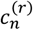 denotes *r*-th graph common representation of node *v*_*n*_, *s*_*n*_ denotes shared variable of node *v*_*n*_.
2. **The correlation loss**. To encourage statistical independence between common and private representations, thereby reducing redundancy and promoting their separation across modalities, SpaMOAL introduces a correlation loss. For each modality *r*, we implement two learnable projection functions *ϕ*^(*r*)^ and *ψ*^(*r*)^ using MLPs with distinct parameters, project the common representation *C*^(*r*)^ and the private representation *P*^(*r*)^ into a comparable latent space, where their dependency is minimized using the Pearson correlation. The correlation loss is defined as:

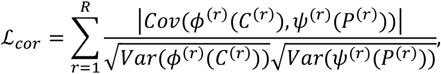

where *Cov*(⋅,⋅) and *Var*(⋅) indicate covariance and variance operations respectively.
3. **The reconstruction loss**. To ensure that both node features and local topological structure are faithfully reconstructed from the learned latent representations, the common and private representations are concatenated and passed through a reconstruction network to generate the reconstructed node representations 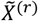. The reconstruction loss is formulated as:

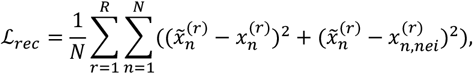

where 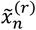 denotes the reconstructed feature vector of node *v_n_*, 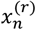 is the original input feature and 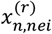 represents the average feature of node *v*_*n*_’s k nearest neighbors in the spatial graph. The first term enforces accurate node feature reconstruction, and the second term preserves neighborhood consistency in the spatial graph.

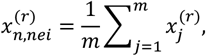

for computational efficiency, we sampled *m* neighbors for each node *v*_*n*_, 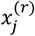 represents the original feature of node *v*_*j*_, which is a 1-hop neighborhood of node *v*_*n*_.
4. **The contrastive loss**. To refine private/modality-specific representations and enhance discriminative power, SpaMOAL applies contrastive learning guided jointly by pseudo-cluster labels derived from group clustering and spatial adjacency. Pseudo-cluster labels provide weak supervision within each private space, while spatial adjacency in the graph constrains the selection of positive and negative pairs to local microenvironments. Positive pairs, denoted as 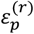, are defined as spatially adjacent spots (i.e., *A_ij_* = 1) that assigned to the same group (*k*_*i*_ = *k*_*j*_), Negative pairs, denoted as 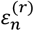, are defined as adjacent spots assigned to different groups (*k*_*i*_ ≠ *k*_*j*_). All other cases are excluded from pair construction.

The contrastive objective encourages neighboring spots belonging to the same group to remain close (i.e., positive pairs), while pushing apart those from different groups (i.e. negative pairs). This spatially constrained optimization emphasizes microenvironment-level coherence and boundary discrimination, enabling each modality to capture locally discriminative patterns. Formally, the contrastive loss is defined as:

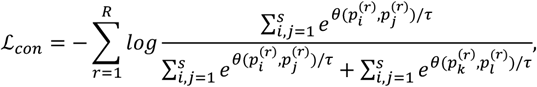

where 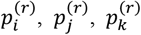 and 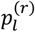 indicate the private representations of node *v*_*i*_, *v*_*j*_, *v*_*k*_ and 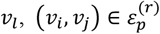 and 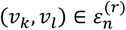, s is the number of sampled 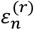, *θ* is the cosine similarity operation and *τ* is a temperature parameter that adjusts the smoothness of similarity scores to control the model’s ability to distinguish between positive and negative samples.

#### Training objective of SpaMOAL

The overall objective function of SpaMOAL is formulated as:

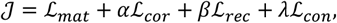

where *α, β* and *λ* are non-negative weighting parameters. Based on empirical validation across diverse datasets, parameter is empirically recommended as *α* ∈ [0.5,1.0], *β* ∈ [0.1,1.0] and *λ* ∈ [0.5,1.0]. A parameter sensitivity analysis for the above parameters has been conducted, with relevant results provided in Supplementary Fig. S1. All private representations *P* are concatenated with the shared variable *S* to form the final representations *Z*. The resulting representations are clustered using mclust^25^, with Leiden and Louvain provided as alternative choices^26^. For all other parameters, we use the default values provided in the code.

### Benchmark datasets

We benchmarked SpaMOAL against two recent representative methods—SpatialGlue and MISO—across 11 datasets comprising 17 samples (Supplementary Table S1). These datasets include: **(i) seven simulated datasets** generated using scMultiSim^27^ or non-negative spatial factorization methods following the procedure by Townes et al.^28^, spanning diverse biological scenarios. Each dataset comprises 500 –12,965 cells and covers two to three omics modalities (e.g., RNA, ATAC, protein). Three datasets also include synthetic H&E-like images with added noise to provide morphological modality; **(ii) a mouse brain development dataset** profiled by MISAR-seq^29^, which jointly measures spatially resolved chromatin accessibility and transcriptome. Four tissue sections spanning different developmental stages (E11, E13.5, E15.5 and E18.5) were analyzed, each encompassing 1,263 to 2,129 spots with varying expert annotations; **(iii) a juvenile mouse brain dataset** comprising a postnatal day 22 (P22) coronal section, profiled for the active promoter mark H3K4me3, enhancer mark H3K27ac, histone modification mark H3K27me3, and ATAC^3^, encompassing 9,215 to 9,752 pixels and annotated with 16 anatomical regions from the Allen Brain Atlas; **(iv) an embryonic day 13 (E13) mouse brain dataset** generated by spatial co-assay ATAC-RNA-seq, containing 2,187 pixels^3^; **(v) a 10X Visium human breast cancer dataset** covering spatial transcriptomics, spatial proteomics (with 35 protein markers), and histological imaging, containing 4,169 spots^30^. Clustering performance on the simulated and mouse brain development datasets (which have gold-standard annotations) were assessed through visualizations in addition to six quantitative metrics: the Adjusted Rand Index (ARI), Adjusted Mutual Information (AMI), Normalized Mutual Information (NMI), Homogeneity, Mutual Information (MI), and V-measure. For datasets lacking expert-annotated labels, the evaluations were based on marker-based and functional biological validation.

## Results

### SpaMOAL achieves state-of-the-art performance across diverse multi-omics simulated data

To quantitatively assess SpaMOAL’s performance, we benchmarked it against SpatialGlue and MISO on seven simulated datasets with known ground-truth labels, including RNA-ATAC dual-omics (Simulations 1, 4, and 6), RNA-ATAC-protein triple-omics (Simulation 2), and image-integrated RNA-ATAC multi-omics (Simulations 3, 5, and 7), each representing distinct yet complementary facets of the ground truth. The RNA-ATAC dual-omics datasets were generated using scMultiSim^27^. To simulate morphological image data, we added random noise to the ground truth annotations to mimic the visual characteristics of H&E-staining. The RNA-ATAC-protein triple-omics dataset was generated following the approach implemented in SpatialGlue.

In the RNA-ATAC dual-omics simulation (Simulation 1), SpaMOAL successfully identified all four simulated spatial domains with high structural continuity in the visualization results. However, SpatialGlue incorrectly identifies both domain1 and domain4. MISO produced fragmented points within domain1 and domain2. And it failed to distinguish the spatially continuous domain3 and domain4 (Fig. 2A-2B). In the RNA-ATAC-protein triple-omics simulation (Simulation 2), both SpaMOAL and SpatialGlue accurately resolved four spatial domains along with a technical noise background. In contrast, MISO incorrectly fragmented the spatially coherent distributed domain1 into scattered spots which is likely because of the absence of explicit spatial constraints in its model architecture (Fig. 2C-2D). In the H&E-integrated dual-omics simulation (Simulation 3) (Fig. 2E), SpaMOAL precisely identified all four spatial domains and their boundaries. SpatialGlue correctly delineated the narrow domain3 and its borders, but failed to fully separate domain4 and domain1. MISO only accurately identified domain2, while the remaining three domains showed substantial deviations in both localization and boundary definition compared to the ground truth. Quantitative metrics confirmed SpaMOAL outperformed MISO and SpatialGlue across all evaluation metrics (Fig. 2F).

**Fig. 2.**
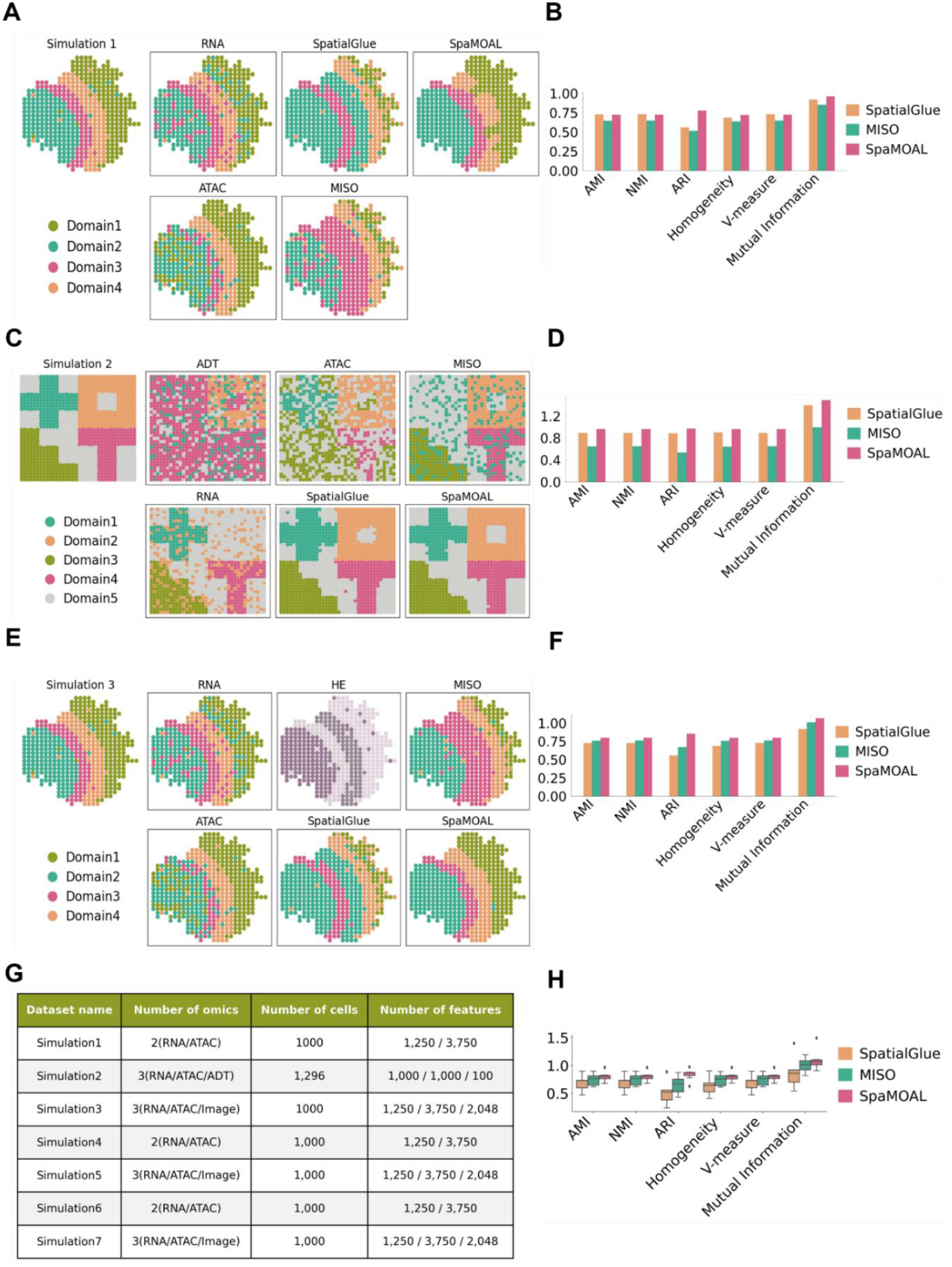
SpaMOAL accurately identifies spatial domains in simulated multi-omics data. A, Spatial plots of Simulation1, which integrates RNA modality and ATAC modality. B, Quantitative evaluation results of methods based on six supervised metrics for Simulation1. C, Spatial plots of Simulation2, which integrates Protein modality, RNA modality and ATAC modality. D, Quantitative evaluation results of methods for Simulation2. E, Spatial plots of Simulation3, which integrates RNA modality, ATAC modality and H&E images. F, Quantitative evaluation results of methods for Simulation3. G, Information statistics (including modality composition, dataset size, feature number) of all seven simulated datasets. H, Box plots of six supervised metrics for three methods across seven simulated datasets.

To further assess robustness, four additional simulated datasets with varied parameters were generated, the visualization results are in Supplementary Fig. S2. The statistics of all seven simulated datasets were summarized in Fig. 2G. Quantitative assessments using the same metrics demonstrated that SpaMOAL consistently achieved superior accuracy and stability (Fig. 2H, Supplementary Table S2).

### SpaMOAL accurately delineates spatial domains in the development of mouse brain tissues profiled by MISAR-seq

To assess the ability of SpaMOAL to resolve fine-grained spatial domains within complex and dynamically developing tissues, we compared it with two representative methods on a multimodal spatial dataset of the developing mouse fetal brain generated by MISAR-seq. MISAR-seq is an advanced spatial multi-omics sequencing platform that jointly profiles high-quality spatial transcriptomics (RNA-seq) and chromatin accessibility landscapes (ATAC-seq) from the same tissue sections. We analyzed four embryonic stages, including E11.0, E13.5, E15.5, and E18.5, to capture a broad spectrum of neurodevelopmental progression. For each stage, we systematically evaluated clustering performance using individual modalities (RNA or ATAC), and further compared the performance of SpaMOAL against SpatialGlue and MISO. SpatialGlue, which does not support image input, was evaluated only without H&E images, whereas SpaMOAL and MISO were evaluated under two settings (with and without H&E images). Expert-curated spatial domain annotations from the original study were served as the gold standard for evaluation (Fig. 3A, 3C and 3E, Supplementary Fig. S3A).

**Fig. 3.**
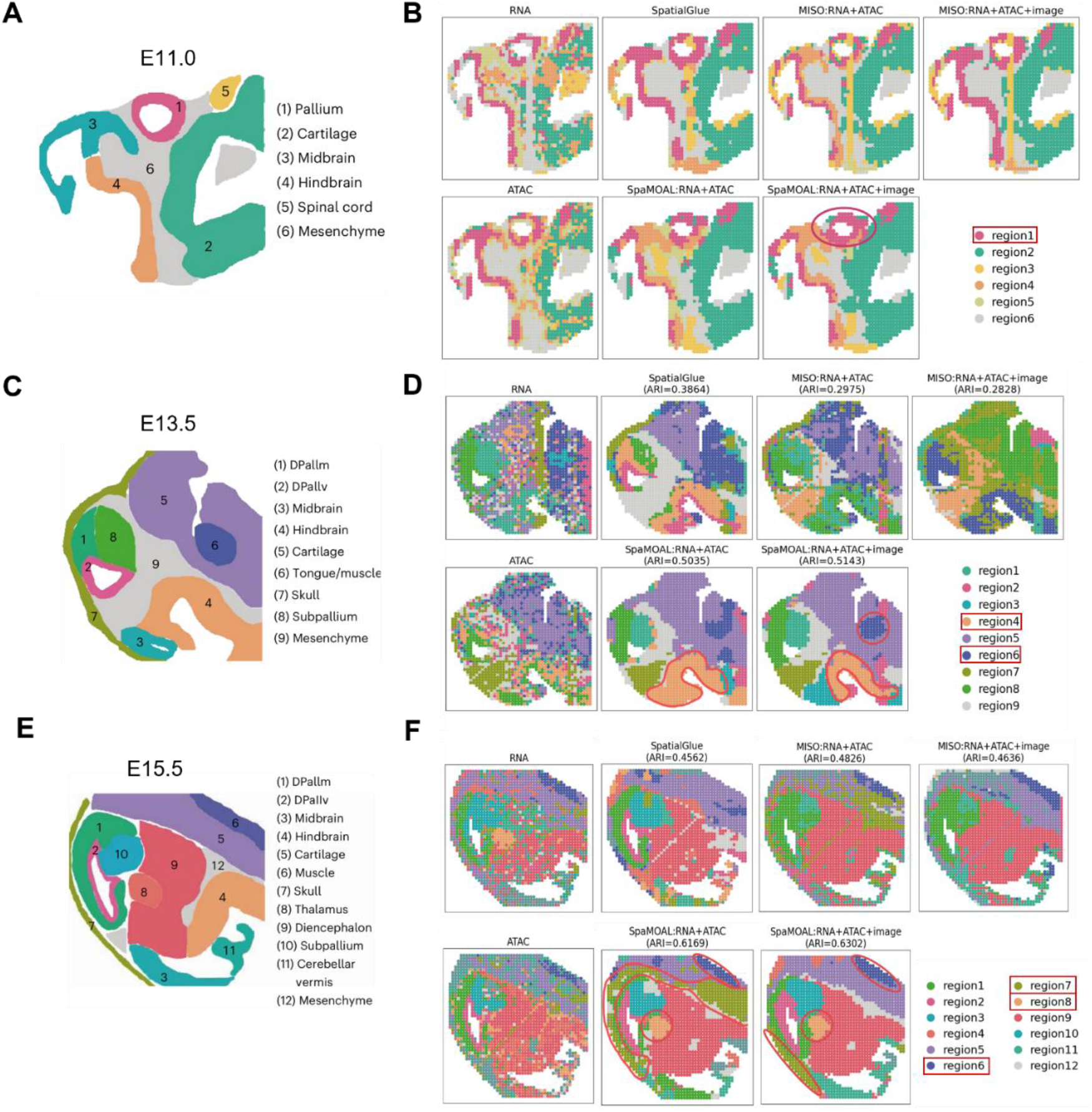
SpaMOAL accurately delineates spatial domains during mouse brain development using MISAR-seq. A, Anatomic annotation of major tissue regions based on the H&E images for E11.0 mouse brain. B, Spatial plots of the E11.0 mouse brain with unimodal clustering (left) and clustering results (right) from spatial multi-omics integration methods—SpatialGlue, MISO and SpaMOAL. C, Anatomic annotation of major tissue regions based on the H&E images for E13.5 mouse brain. D, Spatial plots of the E13.5 mouse brain. E, Anatomic annotation of major tissue regions based on the H&E images for E15.5 mouse brain. F, Spatial plots of the E15.5 mouse brain.

Visual inspection of clustering outcomes indicated that single-omics analyses, based on either RNA or ATAC data alone, failed to capture the coherent spatial organization of the developing mouse brain, producing fragmented and disordered domains. At the E11.0 stage, both SpaMOAL (with image) and SpatialGlue accurately identified the Pallium region as a continuous ring-like structure (region 1), in alignment with known neuroanatomical features. Notably, only SpaMOAL (with and without image) was able to resolve the Cartilage domain as a smooth and spatially coherent structure (region 2). Whereas MISO misassigned the Cartilage domain, producing fragmented and discontinuous partitions (Fig. 3B).

At the E13.5 stage, SpaMOAL (with image) uniquely recovered the Tongue/muscle domain (region 6) as a distinct spatial domain, while all other methods failed to resolve this region (Fig. 3D). Incorporation of image features enabled SpaMOAL to further refine the previously merged region (identified as region 4 without image) encompassing the hindbrain and midbrain into hindbrain (region 4) and midbrain (region 3), suggesting that high-resolution morphological cues provide additional spatial constraints that enhance the fidelity of domain delineation. In contrast, MISO— regardless of image integration—consistently produced fragmented and mosaic-like spurious domains, obscuring the underlying true anatomical structures.

At the later E15.5 stage, SpaMOAL (with or without image) and SpatialGlue accurately recovered canonical neurodevelopmental structures, including the DPallm (region 1), DPallv (region 2), and Subpallium (region 10). Both SpaMOAL (with or without image) and MISO (with image) faithfully delineated the regularly arranged Muscle domain (region 6). Notably, at this stage, SpaMOAL, independent of image usage, was the only method capable of correctly identifying the Skull (region 7) and Thalamus (region 8) as discrete and spatially localized domains, whereas all other methods failed to reconstruct these regions (Fig. 3F). Integration of image information, SpaMOAL further allowedto refine the boundaries of morphologically ambiguous tissues, such as Cartilage (region 5) and Mesenchyme (region 12). These observations indicated that such tissue structures might rely more heavily on morphological cues, thereby giving SpaMOAL’s triple-modal integration (RNA, ATAC, and H&E image) a clear advantage in resolving their spatial architecture. Results of E18.5 are shown in Supplementary Fig. S3B, and quantitative metrics for each stage are provided in Supplementary Fig. S3C-3F.

Taken together across developmental stages, the quantitative metrics further supported the visualization results, with SpaMOAL consistently outperforming MISO and SpatialGlue (Supplementary Fig. S3G). SpaMOAL achieved the closest agreement with expert anatomical annotations, underscoring its superior performance in multimodal data integration and in resolving complex tissue architectures.

### SpaMOAL accurately identifies spatial domains in mouse brain and embryonic tissue samples (spatial ATAC-RNA-seq)

To further demonstrate SpaMOAL’s capability to resolve complex spatial domains in higher resolution spatial multi-omics data, we applied it to mouse brain datasets jointly profiling transcriptomic and epigenomic features. Specifically, we analyzed a postnatal day 22 (P22) mouse brain coronal section profiled using spatial ATAC-RNA-seq, which simultaneously measures mRNA expression and chromatin accessibility within a single tissue slice. This dataset included four sample types: (1) RNA-seq combined with CUT&Tag-seq targeting the active promoter mark H3K4me3; (2) RNA-seq combined with CUT&Tag-seq targeting the enhancer mark H3K27ac; (3) RNA-seq combined with CUT&Tag-seq targeting the repressive mark H3K9me3; and (4) RNA-seq combined with ATAC-seq. To assess the correspondence between SpaMOAL-inferred spatial domains and true anatomical structures, we used high-resolution annotations from the Allen Brain Atlas as the reference. These annotations covered major brain regions, including cortex layers (ctx), genu of the corpus callosum (ccg), lateral septal nucleus (ls), nucleus accumbens (acb), ventrolateral thalamic nucleus (vl), anterior commissure (aco), and caudate putamen (cp) (Fig. 4A). In this benchmark, we systematically evaluated the performance of SpaMOAL against unimodal baselines (RNA-only and ATAC-only) and existing multi-omics approaches (SpatialGlue and MISO).

**Fig. 4.**
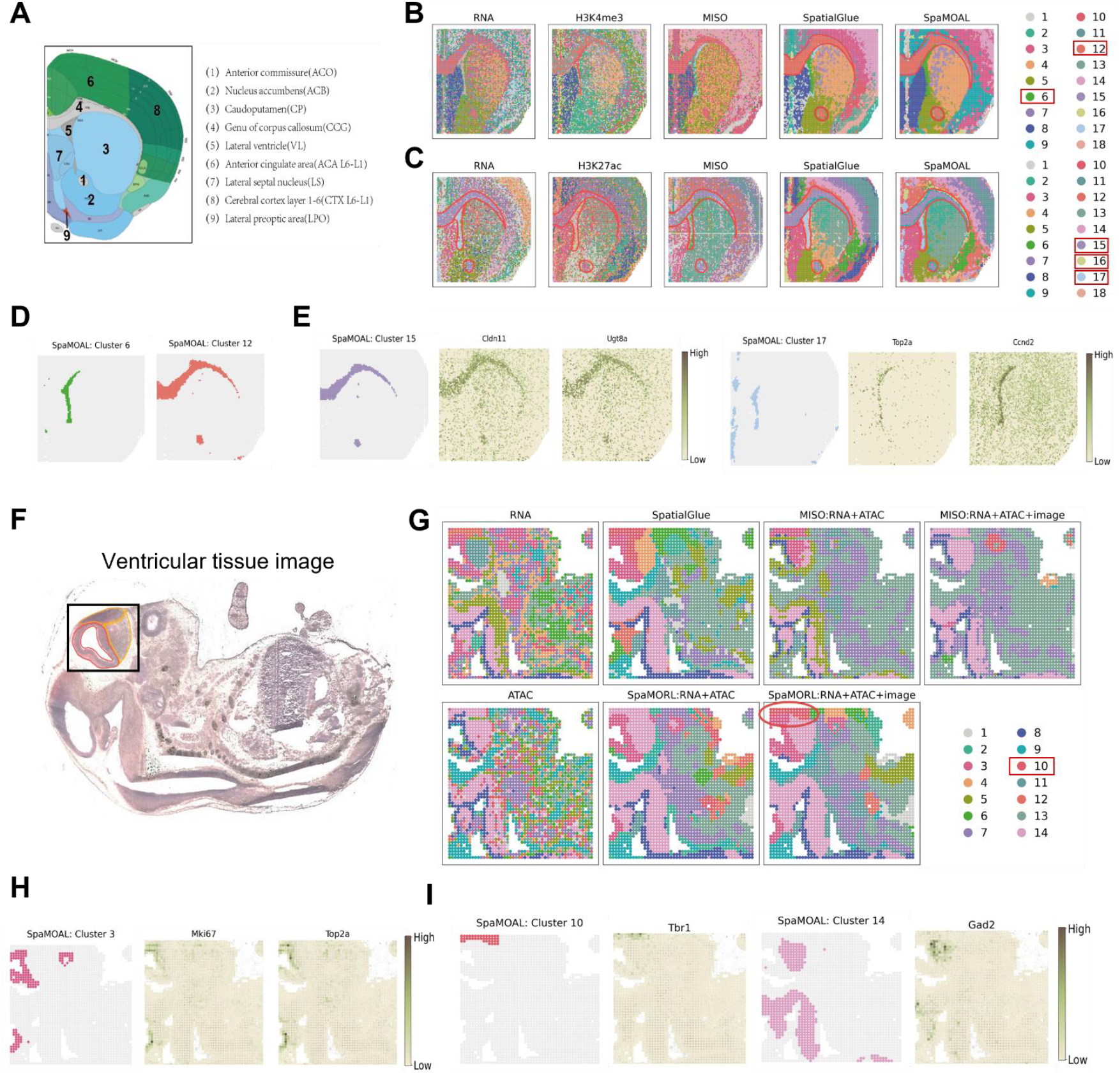
SpaMOAL accurately identified spatial domains in mouse brain samples (RNA-seq and CUT&Tag-seq) and mouse embryonic tissues (RNA-seq and ATAC-seq). A, Annotated reference of the mouse brain coronal section from the Allen Mouse Brain Atlas. B, Spatial plots of RNA and H3K4me3 with unimodal clustering and clustering results from spatial multi-omics integration methods—SpatialGlue, MISO and SpaMOAL. C, Spatial plots of RNA and H3K27ac dataset. D, Separate spatial plots of cluster 6 and cluster 12 identified by SpaMOAL in the mouse brain P22 sample (spatial-CUT&Tag-RNA-seq, H3K4me3). E, Spatial plots of cluster 15 and heatmaps of its marker genes *Cldn11* and *Ugt8a*, along with spatial plots of cluster 17 and its marker genes *Top2a* and *Ccnd2*. F, H&E-stained histology image of an adjacent tissue section. G, Spatial plots of mouse embryo with unimodal clustering(left) and clustering results(right) from spatial multi-omics integration methods—SpatialGlue, MISO and SpaMOAL. H, The Ventricular zone spatial plots of clusters identified by SpaMOAL in the mouse embryo, and spatial gene expression plots of marker genes for the Ventricular zone. I, Separate spatial plots of cluster 3, cluster 10 and cluster 14 identified by SpaMOAL in the mouse embryo.

Across the global visualizations of sample 1 (Fig. 4B) and sample 2 (Fig. 4C), we observed clear differences among methods in their ability to recover spatial organization. Unimodal approaches (RNA-only or ATAC-only), which lack complementary molecular signals and explicit spatial constraints, tend to produce more fragmented and spatially dispersed clusters that fail to delineate continuous anatomical structures. Although MISO leverages multi-omics signals and could recover certain regions (e.g., the ccg), it does not explicitly model spatial coordinates, resulting in fragmented output and numerous spurious micro-clusters. In contrast, SpaMOAL exhibits markedly improved spatial coherence, yielding smoother domain boundaries and more continuous structures.

Specifically, in sample 1, all three multi-omics integration methods, SpaMOAL, MISO, and SpatialGlue, consistently identified the aco and the ccg (cluster 12), indicating that these regions exhibit robust molecular signatures across methods. Notably, SpaMOAL and SpatialGlue were able to accurately resolve and fully delineate the vl (cluster 6), a narrow laminar structure situated between the ccg, ls, and cp. This region functions as a critical relay station in the motor thalamo-cortical circuit, contributing to motor initiation, temporal coordination, and parkinsonian bradykinesia. By contrast, MISO offered an incomplete depiction of this fine-grained structure, underscoring their limitations in resolving regions with weak signals or diffuse boundaries (Fig. 4B and 4D). In sample 2, consistent with the results from sample1, SpaMOAL, MISO, and SpatialGlue consistently detected the aco and ccg (cluster 15), whereas accurate resolution and complete delineation of the vl (cluster 17) were achieved only by SpaMOAL and SpatialGlue (Fig. 4C). SpaMOAL uniquely identified a thin outer ctx (cluster 16) at the periphery of the ccg. This region exhibits weak signal intensity, sparse cellularity, and subtle laminar morphology, corresponding to an anatomically elusive marginal layer (Fig. 4E). The other two methods failed to resolve this structure, indicating their limited sensitivity to weak chromatin accessibility or low-abundance transcriptional signals. Together, these results demonstrated that SpaMOAL achieves superior resolution and spatial coherence in dissecting complex brain architectures, enabling not only the identification of canonical anatomical domains but also the discovery of morphologically cryptic and transcriptionally subtle laminar structures. Results of sample 3 and sample 4 were presented in Supplementary Fig. S4.

We next extended our analysis to a spatial ATAC-RNA-seq dataset derived from embryonic day 13 (E13) mouse brain sections (Fig. 4F). Visualization results revealed that SpaMOAL (with or without image integration) accurately delineated the fine substructure of the ventricular zone (cluster 3) within the telencephalon, demonstrating strong capacity for spatial multi-omics resolution (Fig. 4G). Anatomically, the ventricular zone is primarily composed of neural stem and progenitor cells, with canonical markers such as *MKI67* and *TOP2A* being highly enriched in neural progenitors^31^. Our spatial partitioning results validated by these molecular features: the high expression of *MKI67* and *TOP2A* exhibited pronounced spatial colocalization with cluster 3, thereby confirming its correspondence to neural progenitor populations within the ventricular zone (Fig. 4H). Notably, upon integration of histological images, SpaMOAL exhibited markedly improved in resolution and accuracy, clearly distinguishing the three major telencephalic subregions-ventricular zone (cluster 3), subpallium (cluster 14), and pallium (cluster 10) (Fig. 4I). This observation underscores the advantage of SpaMOAL in image-guided integration and highlights its potential to achieve high-precision, multimodal spatial characterization in complex developing tissues.

### SpaMOAL accurately identified spatial domains in human breast cancer tissue (spatial RNA-Protein-seq)

Spatial multi-omics technologies provide a comprehensive framework for systematically resolving tissue organization across molecular and spatial dimensions. These approaches elucidate normal cellular organization during development and uncover disrupted tissue architecture in disease, particularly in cancer. Within the tumor immune microenvironment, tertiary lymphoid structures (TLSs) represent a critical immunological niche that supports antigen presentation, lymphocyte activation, and the maintenance of anti-tumor immune responses. The presence and maturation state of TLSs are strongly linked to improved patient survival in untreated tumors, underscoring the need for accurate spatial identification and characterization of TLSs to better understand tumor immunity and guide prognostic assessment. To evaluate the capacity of SpaMOAL to resolve complex immune microenvironments in human tissues, we applied it to spatial datasets derived from human breast cancer sections, comprising spatial transcriptomics, spatial proteomics (35 protein markers), and histological imaging (Fig. 5A). We additionally benchmarked SpaMOAL against unimodal RNA- or protein-based methods, as well as existing multimodal integration approaches (SpatialGlue and MISO), to systematically assess its performance in resolving biologically coherent spatial domains (Fig. 5B).

**Fig. 5.**
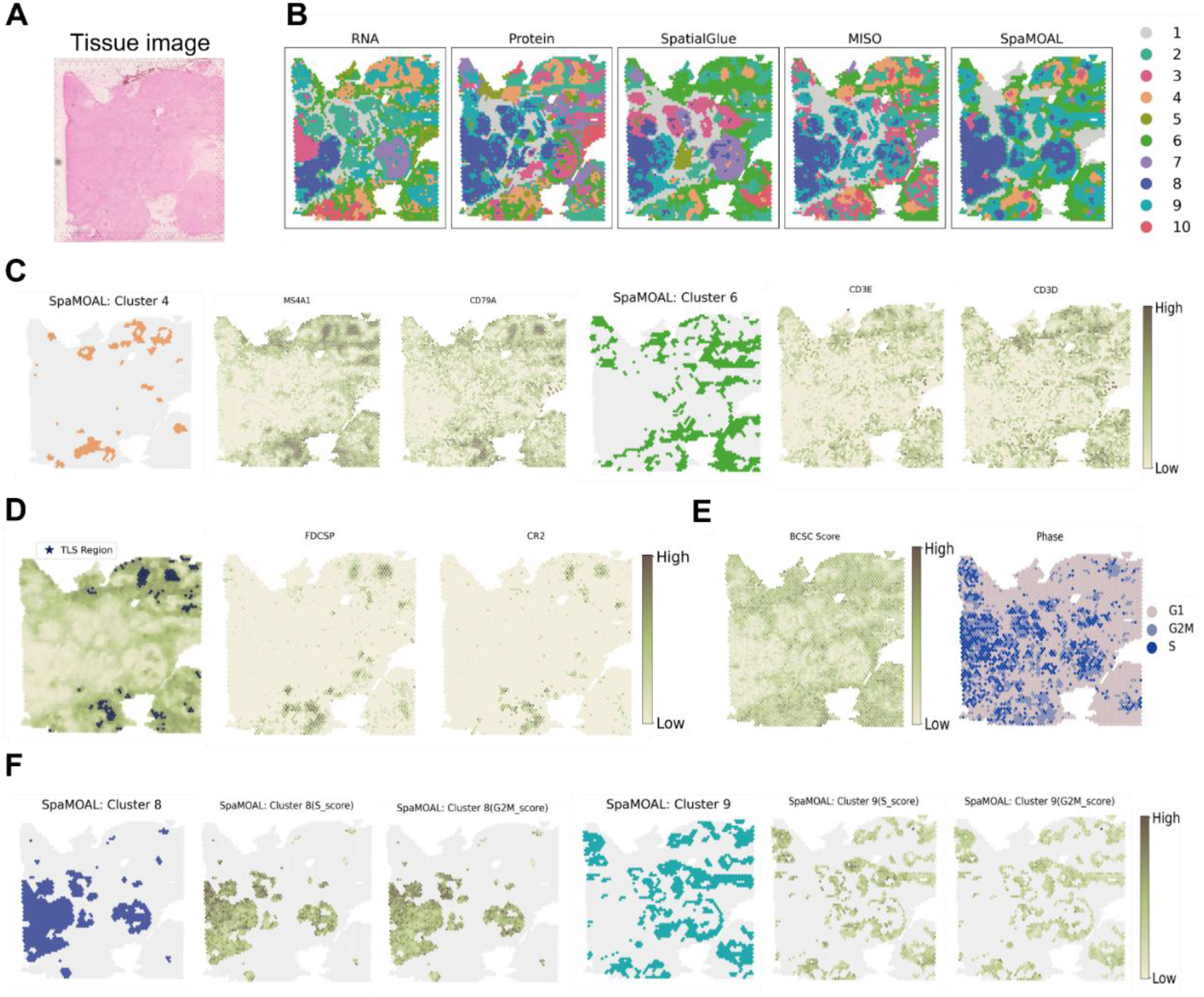
SpaMOAL accurately identified spatial domains in human breast cancer tissue (spatial RNA-Protein-seq). A, Tissue image of human breast cancer dataset. B, Spatial plots of RNA and Protein with unimodal clustering and clustering results from spatial multi-omics integration methods—SpatialGlue, MISO and SpaMOAL. C, The spatial plots of cluster 4 and cluster 6 identified by SpaMOAL, and spatial gene expression plots of marker genes for the clusters. D, TLS score and spatial gene expression plots of TLS marker genes. E, Stemness score plots. F, Separate spatial plots of cluster 8 and cluster 9 identified by SpaMOAL, and their cell-cycle score plots.

In the resulting clustering analyses, both SpaMOAL and MISO robustly identified TLS regions in human breast cancer samples, and consistently annotated cluster 4 and cluster 6 as TLS-enriched compartments. Within TLS region, cluster 4 corresponds to the TLS core enriched with prominent B cell accumulation, which is validated by the high expression of *MS4A1* and *CD79A*. In contrast, cluster 6 represents the surrounding T cell-enriched TLS peripheral zone, the positive expression of *CD3E* and *CD3D* further confirms the cellular composition of this region (Fig. 5C). The distribution pattern reflects the typical zonal architectural features of mature TLS. To delineate TLS regions with high precision in spatial transcriptomic data, we performed spatial annotation using a curated set of 50 TLS-associated signature genes compiled from previous studies^32^. Classical TLS marker genes, including *LTB* and *TMSB4X*, were markedly upregulated within these regions (Fig. 5D), demonstrating SpaMOAL’s capability to precisely capture the specific molecular signature of TLSs.

We further quantified breast cancer stemness by computing a stemness score based on six canonical markers (*CD44, ALDH1A1, ALDH1A3, PROM1, SOX2, CXCR4*) (Fig. 5E). Notably, SpaMOAL uniquely resolved the basal-like tumor compartment into two functionally distinct subregions: a proliferative region (cluster 8) enriched for S/G2M phase signatures and a quiescent region (cluster 9) characterized by low cell-cycle activity^33^. This was not explicitly distinguished by other multimodal integration approaches. Despite the fact that this region exhibited relatively weak transcriptional signals, spatial patterns were clearly displayed at the protein level and in tissue morphology. These findings suggest that SpaMOAL effectively integrates proteomic and histological cues with high fidelity, enabling the detection of biologically meaningful tumor subregions with minimal transcriptomic contrast. To further assess cellular functional states, we applied scFates^34^ to infer pseudotime trajectories across spatial location. SpaMOAL-derived cluster 8 exhibited markedly elevated pseudotime scores, consistent with enrichment of proliferative G2/M and S phases (Supplementary Fig. S5). In contrast, SpaMOAL-derived cluster 9 showed relatively lower pseudotime values, with most cells residing in G1 or quiescent states (Fig. 5F). These findings not only highlight the spatial heterogeneity of cell-cycle dynamics within the tumor microenvironment but also demonstrate SpaMOAL’s capability to resolve tumor subpopulations with distinct proliferative potentials.

Collectively, by capturing spatially coherent and functionally relevant tissue domains defined jointly by protein expression and morphological features—rather than relying solely on transcript-level variation—SpaMOAL achieves sensitive and high-resolution characterization of dynamic cellular states through integrated multimodal profiling.

## Discussion

The rapid development of spatial multi-omics technologies has created new opportunities for accurately delineating complex spatial tissue architectures, which is critical for understanding cellular function and tissue organization under both physiological and pathological conditions. Here, we introduce SpaMOAL, a graph contrastive autoencoder-based method, for integrative spatial domain detection that combines spatial coordinates, molecular profiles, and histological images into unified multimodal representations. We systematically benchmarked SpaMOAL across 17 samples spanning 11 datasets, comparing its performance against single-modality baselines and two state-of-the-art multi-omics integration methods. Across multiple quantitative metrics and visualizations, SpaMOAL consistently outperformed competing methods. From simulated datasets and developing mouse brain to human cancer multi-omics data, the spatial domains identified by SpaMOAL exhibited higher concordance with known biological structures, demonstrating its robustness and broad applicability.

In addition to improved accuracy, SpaMOAL also achieved competitive computational efficiency. Across datasets of increasing complexity, SpaMOAL maintained substantially lower running times than MISO—particularly on the mouse brain RNA-ATAC dataset, where MISO required over 1,800 seconds while SpaMOAL completed analysis in under 1,000 seconds—while performing comparably to or faster than SpatialGlue (Supplementary Fig. S6). These results highlight SpaMOAL’s computational scalability for large multimodal spatial datasets.

While SpaMOAL demonstrates strong overall performance, several aspects warrant further refinement. Although it conceptually separates shared and modality-specific representations, this disentanglement remains partial, as optimal performance is achieved using fused embeddings. Additionally, despite its integrative capability, the interpretability of SpaMOAL is limited, as the individual contributions of each modality to spatial domain formation cannot yet be quantitatively determined. Addressing these issues through interpretable latent modeling and modality-attribution analysis may provide deeper biological insight. Furthermore, because current spatial multi-omics technologies are restricted to RNA, ATAC, and protein measurements, our ability to fully resolve transcriptional regulatory networks remains constrained. Integrating additional omics layers, such as DNA methylation or histone modifications, could further enhance SpaMOAL’s capacity to map transcriptional networks, offering a more comprehensive view of the underlying regulatory architecture. As spatial multi-omics technologies continue to advance, generating increasingly large and diverse datasets, SpaMOAL is well positioned to leverage these data for time-series or multi-sample analyses, providing new perspectives on spatial organization during tissue development, disease progression, and therapeutic response.

## Data availability

The simulated datasets are available at https://github.com/XiangyuLi-Lab/SpaMOAL. The MISAR-seq mouse brain dataset is accessible at the National Genomics Data Center with accession number OEP003285. The spatial ATAC-RNA-seq mouse brain dataset can be found at https://web.atlasxomics.com/visualization/Fan. Spatial ATAC-RNA-seq mouse embryonic day 13 (E13) data reported in https://cells.ucsc.edu/?ds=brain-spatial-omics. 10x Visium human breast cancer gene and protein expression data can be found at https://www.10xgenomics.com/resources/datasets/gene-and-protein-expression-library-of-human-breast-cancer-cytassist-ffpe-2-standard.

## Code availability

The code of SpaMOAL is available at https://github.com/XiangyuLi-Lab/SpaMOAL.

## Supplementary information for ‘Integrating spatial multi-omics data to decipher spatial domains with SpaMOAL’

### Supplementary Notes

#### 1. Simulation data generation

To provide evaluation data with ground truth, we generated seven multi-modal datasets. Six were created using scMultiSim27, and the remaining one was generated by SpatialGlue19, following the approach outlined by Townes et al.28. Specifically, we first used scMultiSim to generate three RNA-ATAC dual-modal datasets, and then produced three image-integrated RNA-ATAC triple-modal datasets by incorporating simulated H&E images. scMultiSim enables simultaneous modeling of cell identity, gene regulatory networks, cell-cell interactions, and chromatin accessibility, while also introducing technical noise. We generated datasets with different hierarchical structures by varying configurations such as gene regulatory networks, cell numbers, gene counts, and random seeds, detailed configurations are provided in Supplementary Table S3. As scMultiSim does not support the generation of triple-omics data, we directly adopted the RNA-ATAC-protein dataset generated by SpatialGlue. Each data type was modeled using a tailored statistical distribution, the transcriptomic data were modeled with a zero-inflated negative binomial distribution to accurately represent their high sparsity and over-dispersion characteristics. Both proteomic data and chromatin accessibility measurements were modeled using a standard negative binomial distribution, which is appropriate for characterizing their count-based data features.

#### 2. Ablation studies

To evaluate the contribution of each loss function to SpaMOAL’s performance, we conducted ablation studies on the Simulation1 dataset. As shown in Supplementary Table S4, when the model lacks the matching loss (*ℒ*_*mat*_), correlation loss (*ℒ*_*cor*_), reconstruction loss (*ℒ*_*rec*_) or contrastive loss (*ℒ*_*con*_), its performance degrades to varying degrees. This demonstrates the positive contributions of each of the aforementioned components to the model’s performance.

#### 3. Sensitivity to parameters

We evaluated the sensitivity of SpaMOAL to its key hyperparameters—the loss coefficients α, β, and λ—using simulated datasets. The default values were set to α = 0.8, β = 0.5, and λ = 0.9. To assess the effect of varying a single parameter, we held two parameters constant at their defaults while adjusting the third. As shown in Supplementary Fig. S4, although variations in these parameters did influence performance, the overall effect was modest, indicating that SpaMOAL is robust to a range of hyperparameter values.

## Supplementary Figures

**Supplementary Figure S1.**
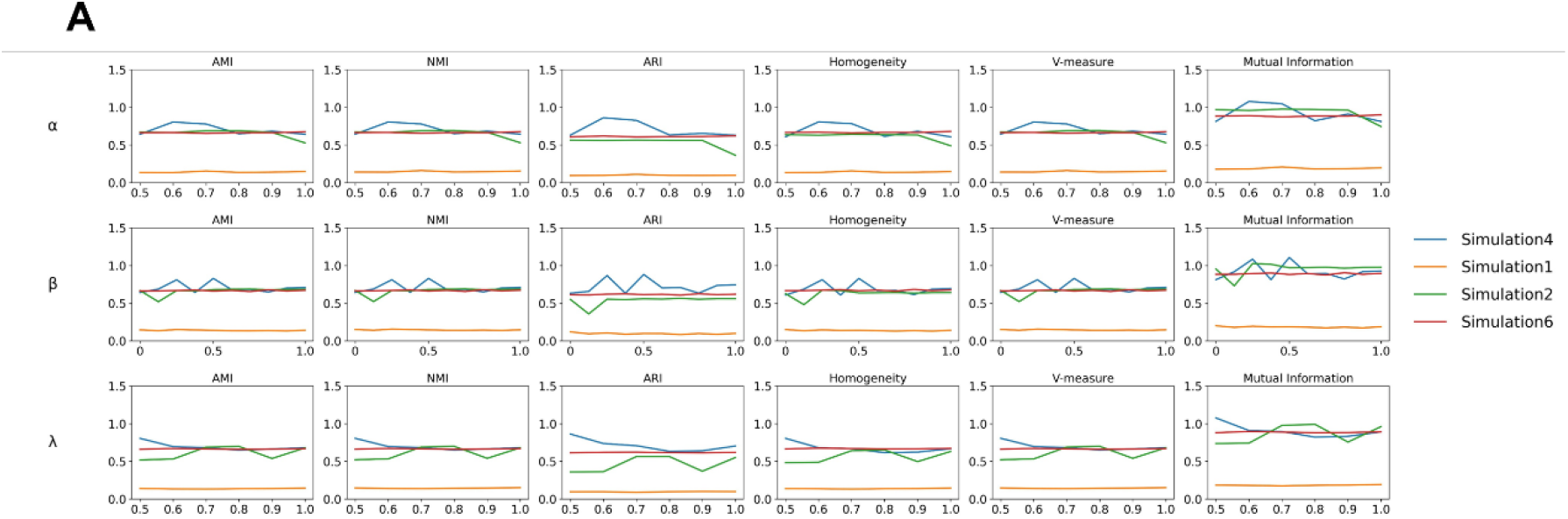
Sensitivity to parameters. A, The line plots illustrate the performance of SpaMOAL of different parameters α, β, and λ, evaluated across four datasets (Simulation 1-4) using six metrics: the Adjusted Rand Index (ARI), Adjusted Mutual Information (AMI), Normalized Mutual Information (NMI), Homogeneity, Mutual Information (MI), and V-measure.

**Supplementary Figure S2.**
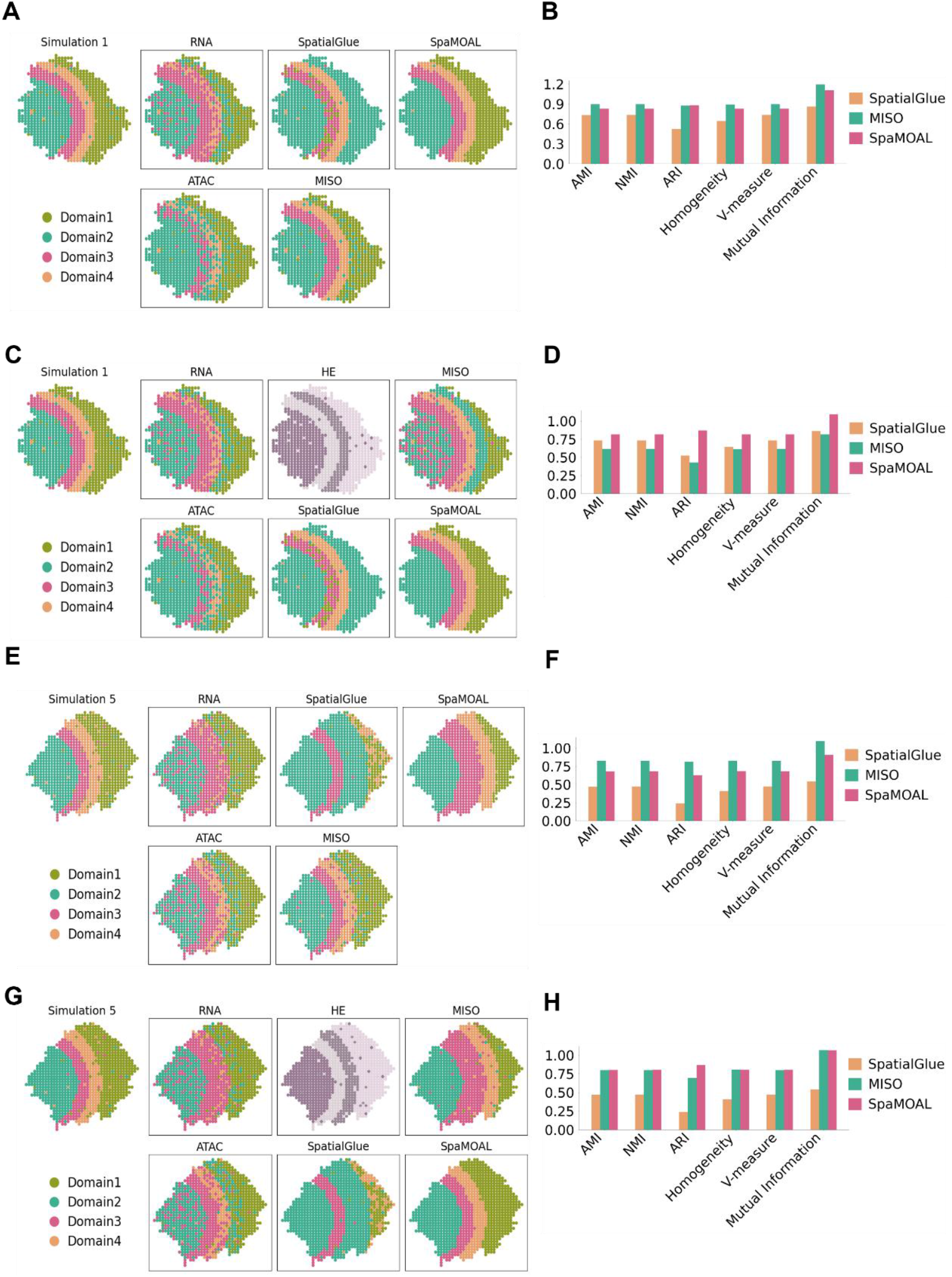
SpaMOAL accurately identifies spatial domains in simulated multi-omics data. A, Spatial plots of Simulation4, which integrates RNA modality and ATAC modality. B, Quantitative evaluation results of methods based on six supervised metrics for Simulation4. C, Spatial plots of Simulation5, which integrates RNA modality, ATAC modality and HE. D, Quantitative evaluation results of methods for Simulation5. E, Spatial plots of Simulation6, which integrates RNA modality and ATAC modality. F, Quantitative evaluation results of methods for Simulation6. G, Spatial plots of Simulation7, which integrates RNA modality, ATAC modality and HE. H, Quantitative evaluation results of methods for Simulation7.

**Supplementary Figure S3.**
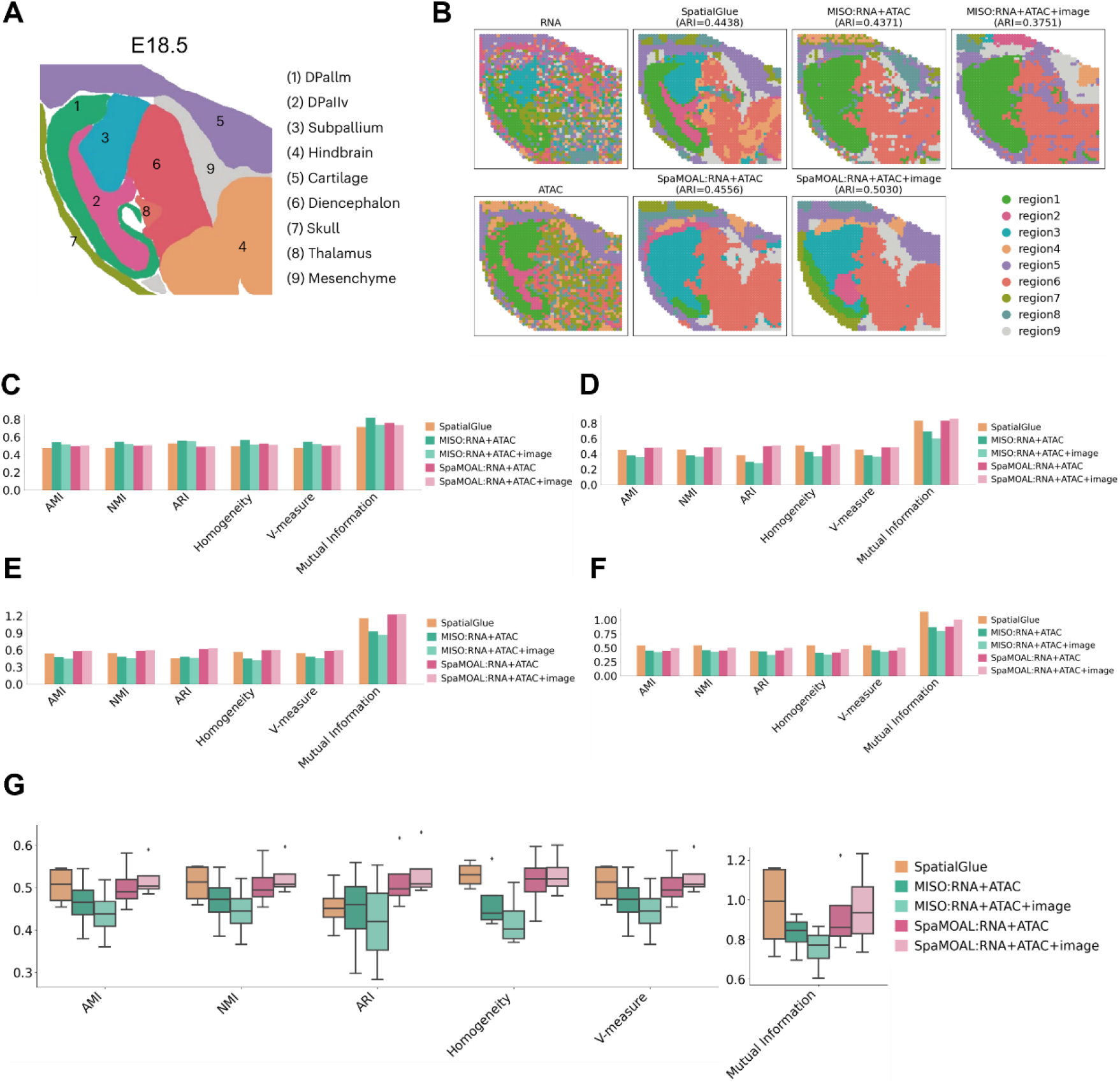
SpaMOAL accurately delineates spatial domains during mouse brain development using MISAR-seq. A, Anatomic annotation of major tissue regions based on the H&E images for E18.5 mouse brain. B, Spatial plots of the E18.5 mouse brain with unimodal clustering (left) and clustering results (right) from spatial multi-omics integration methods—SpatialGlue, MISO and SpaMOAL. C, Quantitative evaluation results of methods based on six supervised metrics for E11.0. D, Quantitative evaluation results of methods for E13.5. E, Quantitative evaluation results of methods for E15.5. F, Quantitative evaluation results of methods for E18.5. G, Box plots of six supervised metrics for three methods across four samples.

**Supplementary Figure S4.**
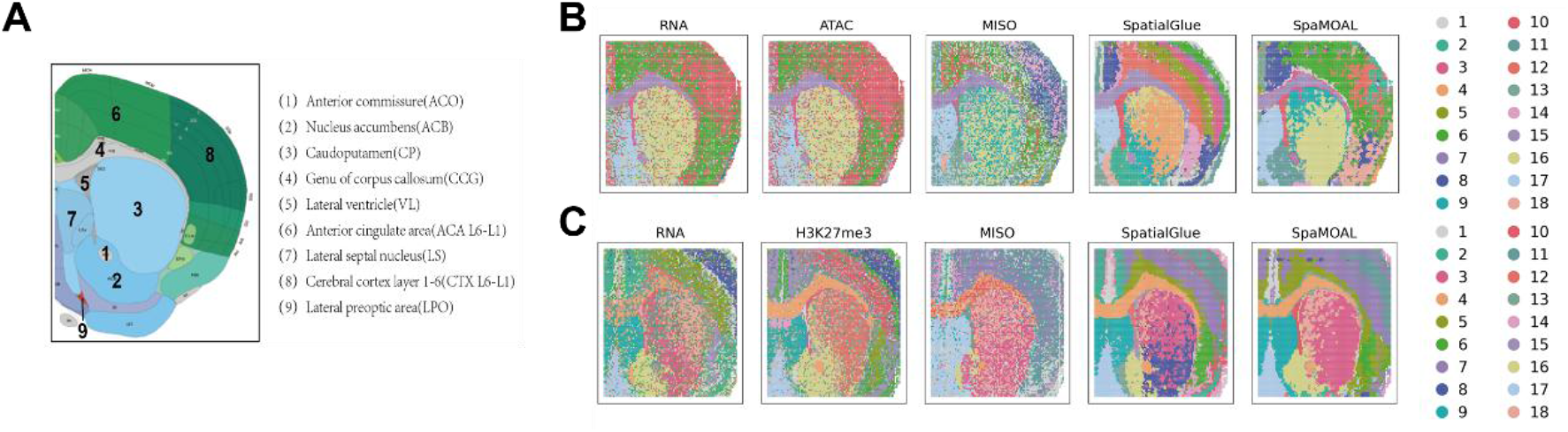
SpaMOAL accurately identified spatial domains in mouse brain samples (RNA-seq and CUT&Tag-seq) and mouse embryonic tissues (RNA-seq and ATAC-seq). A, Annotated reference of the mouse brain coronal section from the Allen Mouse Brain Atlas. B, Spatial plots of RNA and ATAC with unimodal clustering and clustering results from spatial multi-omics integration methods—SpatialGlue, MISO and SpaMOAL. C, Spatial plots of RNA and H3K27me3 dataset.

**Supplementary Figure S5.**
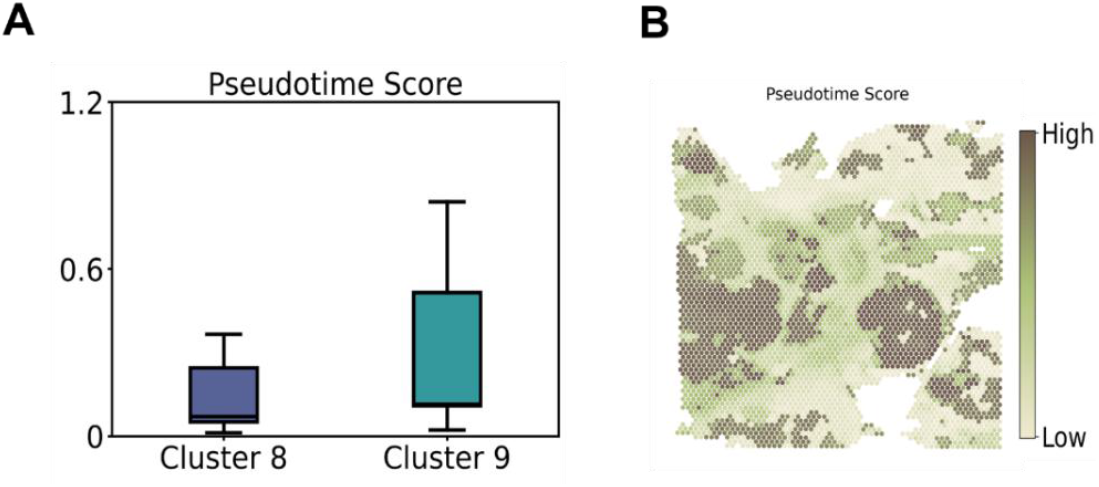
SpaMOAL accurately identified spatial domains in human breast cancer tissue (spatial RNA-Protein-seq). A, Box plots of pseudotime score for cluster 8 and cluster9. B, Spatial pseudotime score heatmap.

**Supplementary Figure S6.**
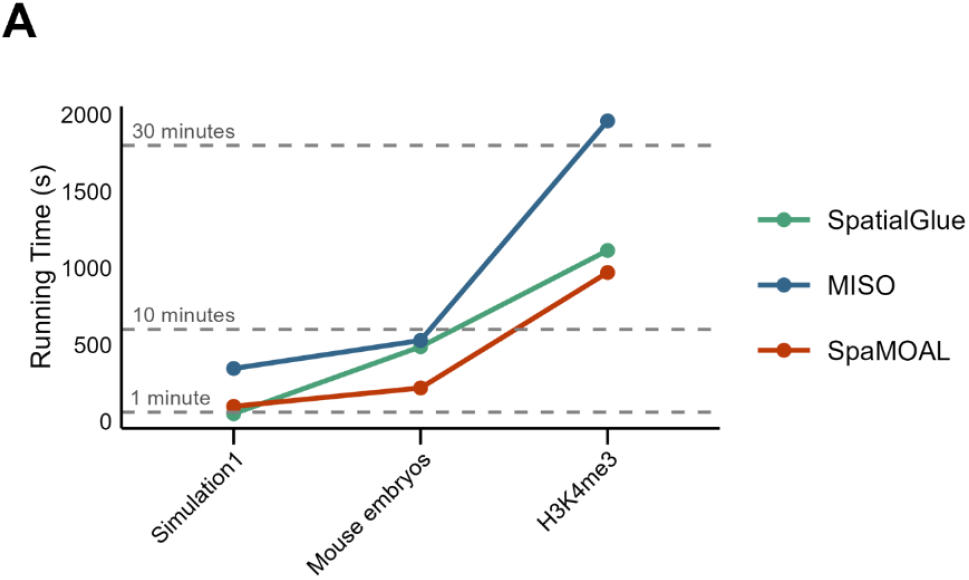
A, Running Time of SpatialGlue, MISO, and SpaMOAL on Three Datasets.

## Supplementary Tables

**Supplementary Table S1.** Experimental datasets used in the manuscript.

| **Dataset** | **Name** | **Platform** | **Size (spots x genes/proteins/peaks/image)** |
| --- | --- | --- | --- |
| **Dataset1** | Simulation1 | scMultiSim | 500x200<br>500x600 |
| **Dataset2** | Simulation2 | NSF (Townes et al., 2023) | 1,296x1,000<br>1,296x100 |
| **Dataset3** | Simulation3 | scMultiSim | 500x200; 500x600<br>500x2,048 |
| **Dataset4** | Simulation4 | scMultiSim | 1,000x1,250<br>1,000x3,750 |
| **Dataset5** | Simulation5 | scMultiSim | 1,000x1,250<br>1,000x3,750<br>1,000x2,048 |
| **Dataset6** | Simulation6 | scMultiSim | 1,000x1,250;<br>1,000x3,750 |
| **Dataset7** | Simulation7 | scMultiSim | 1,000x1,250<br>1,000x3,750<br>1,000x2,048 |
| **Dataset8** | Mouse Brain E11.0 | MISAR-seq | 1,263x31,433<br>1,263x24,333 |
| **Dataset9** | Mouse Brain E13.5 | MISAR-seq | 1,777x31,433<br>1,777x24,333 |
| **Dataset10** | Mouse Brain E15.5 | MISAR-seq | 1,949x31,433<br>1,949x24,333 |
| **Dataset11** | Mouse Brain E18.5 | MISAR-seq | 2,129x31,433<br>2,129x24,333 |
| **Dataset12** | Mouse Brain RNA H3K4me3 | Spatial transcriptome epigenome | 9,548x22,731<br>9,548x35,270 |
| **Dataset13** | Mouse Brain RNA H3K27ac | Spatial transcriptome epigenome | 9,370x23,415<br>9,370x104,162 |
| **Dataset14** | Mouse Brain RNA ATAC P22 | Spatial transcriptome epigenome | 9,215x22,914<br>9,215x121,068 |
| **Dataset15** | Mouse Brain RNA H3K27me3 | Spatial transcriptome epigenome | 9,752x25,881<br>9,752x70,470 |
| **Dataset16** | Mouse Embryo | spatial ATAC-RNA-seq | 2,187x17,058<br>2,187x15,521 |

**Supplementary Table S2.** Quantitative metrics of all methods on simulated datasets.

| Dataset | Method | AMI | NMI | ARI | Homogeneity | V-measure | MI |
| --- | --- | --- | --- | --- | --- | --- | --- |
| Simulation1 | SpatialGlue | 0.7285 | 0.7305 | 0.5595 | 0.6878 | 0.7305 | 0.9196 |
|  | MISO | 0.6458 | 0.6483 | 0.5159 | 0.6363 | 0.6483 | 0.8508 |
|  | SpaMOAL | 0.7223 | 0.7242 | 0.7783 | 0.7189 | 0.7242 | 0.9611 |
| Simulation2 | SpatialGlue | 0.8961 | 0.8965 | 0.8874 | 0.9043 | 0.8965 | 1.3993 |
|  | MISO | 0.6490 | 0.6504 | 0.5374 | 0.6431 | 0.6504 | 0.9952 |
|  | SpaMOAL | 0.9647 | 0.9649 | 0.9748 | 0.9649 | 0.9649 | 1.4930 |
| Simulation3 | SpatialGlue | 0.7285 | 0.7305 | 0.5595 | 0.6878 | 0.7305 | 0.9196 |
|  | MISO | 0.7626 | 0.7642 | 0.6723 | 0.7588 | 0.7642 | 1.0145 |
|  | SpaMOAL | 0.8005 | 0.8019 | 0.8564 | 0.7997 | 0.8019 | 1.0692 |
| Simulation4 | SpatialGlue | 0.7335 | 0.7346 | 0.5260 | 0.6444 | 0.7346 | 0.8614 |
|  | MISO | 0.8962 | 0.8965 | 0.8748 | 0.8930 | 0.8965 | 1.1936 |
|  | SpaMOAL | 0.8279 | 0.8285 | 0.8806 | 0.8273 | 0.8285 | 1.1059 |
| Simulation5 | SpatialGlue | 0.7335 | 0.7346 | 0.5260 | 0.6444 | 0.7346 | 0.8614 |
|  | MISO | 0.6163 | 0.6176 | 0.4292 | 0.6118 | 0.6176 | 0.8178 |
|  | SpaMOAL | 0.8168 | 0.8175 | 0.8697 | 0.8179 | 0.8175 | 1.0933 |
| Simulation6 | SpatialGlue | 0.4696 | 0.4717 | 0.2391 | 0.4090 | 0.4717 | 0.5421 |
|  | MISO | 0.8266 | 0.8271 | 0.8114 | 0.8277 | 0.8271 | 1.0971 |
|  | SpaMOAL | 0.6800 | 0.6811 | 0.6267 | 0.6847 | 0.6811 | 0.9074 |
| Simulation7 | SpatialGlue | 0.4696 | 0.4717 | 0.2391 | 0.4090 | 0.4717 | 0.5421 |
|  | MISO | 0.8012 | 0.8019 | 0.6956 | 0.8062 | 0.8019 | 1.0686 |
|  | SpaMOAL | 0.8039 | 0.8045 | 0.8699 | 0.8045 | 0.8045 | 1.0662 |

**Supplementary Table S3.**
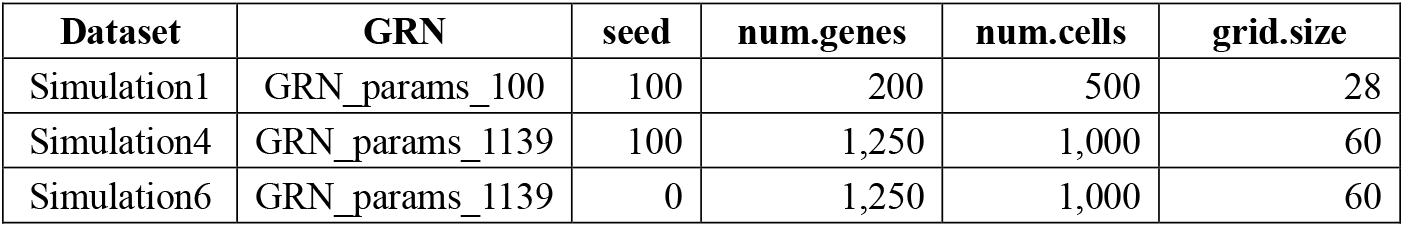
Parameter Configuration for Simulated Datasets.

**Supplementary Table S4.** Results of ablation studies on simulated datasets.

| Method | AMI | NMI | ARI | Homogeneity | V-measure | MI |
| --- | --- | --- | --- | --- | --- | --- |
| **SpaMOAL** | **0.8005** | **0.8019** | **0.8564** | **0.7997** | **0.8019** | **1.0692** |
| w/o $\mathcal{L}_{mat}$ | 0.6785 | 0.6807 | 0.6318 | 0.6776 | 0.6807 | 0.9060 |
| w/o $\mathcal{L}_{cor}$ | 0.6718 | 0.6741 | 0.6271 | 0.6703 | 0.6741 | 0.8962 |
| w/o $\mathcal{L}_{rec}$ | 0.6584 | 0.6607 | 0.6023 | 0.6509 | 0.6607 | 0.8702 |
| w/o $\mathcal{L}_{con}$ | 0.6559 | 0.6582 | 0.6190 | 0.6544 | 0.6582 | 0.8749 |

